# *Shigella* OspF blocks rapid p38-dependent priming of the NAIP–NLRC4 inflammasome

**DOI:** 10.1101/2025.02.01.636075

**Authors:** Elizabeth A. Turcotte, Kyungsub Kim, Kevin D. Eislmayr, Lisa Goers, Patrick S. Mitchell, Cammie F. Lesser, Russell E. Vance

## Abstract

The NAIP–NLRC4 inflammasome senses pathogenic bacteria by recognizing the cytosolic presence of bacterial proteins such as flagellin and type III secretion system (T3SS) subunits. In mice, the NAIP–NLRC4 inflammasome provides robust protection against bacterial pathogens that infect intestinal epithelial cells, including the gastrointestinal pathogen *Shigella flexneri*. By contrast, humans are highly susceptible to *Shigella*, despite the ability of human NAIP–NLRC4 to robustly detect *Shigella* T3SS proteins. Why the NAIP–NLRC4 inflammasome protects mice but not humans against *Shigella* infection remains unclear. We previously found that human THP-1 cells infected with *Shigella* lose responsiveness to NAIP–NLRC4 stimuli, while retaining sensitivity to other inflammasome agonists. Using mT3Sf, a “minimal *Shigella*” system, to express individual secreted *Shigella* effector proteins, we found that the OspF effector specifically suppresses NAIP–NLRC4-dependent cell death during infection. OspF was previously characterized as a phosphothreonine lyase that inactivates p38 and ERK MAP kinases. We found that p38 was critical for rapid priming of NAIP–NLRC4 activity, particularly in cells with low NAIP–NLRC4 expression. Overall, our results provide a mechanism by which *Shigella* evades inflammasome activation in humans, and describe a new mechanism for rapid priming of the NAIP–NLRC4 inflammasome.

## Introduction

Inflammasomes are cytosolic sensors of pathogen ligands, pathogen activities, and other noxious stimuli. Upon triggering, inflammasomes recruit and activate caspases that induce cell death and the release of the proinflammatory cytokines interleukin (IL)-18 and IL-1β. The NAIP–NLRC4 inflammasome is activated upon NAIP recognition of specific bacterial proteins, including flagellin, or the rod or needle proteins of bacterial type III secretion systems (T3SS) (1-5). A single ligand-bound NAIP assembles with multiple copies of NLRC4 (6, 7), which then recruit and activate Caspase-1, a protease that processes pro-IL-18 and pro-IL-1β into their active forms. Caspase-1 also cleaves and activates Gasdermin-D (GSDMD), a pore forming protein that initiates pyroptotic cell death (8, 9). Another inflammatory caspase, Caspase-4, forms a ‘non-canonical’ inflammasome upon direct recognition of cytosolic LPS. Active Caspase-4 also cleaves GSDMD to induce pyroptosis (10, 11). Pyroptosis is important for eliminating infected cells, and for inducing a proinflammatory response to halt infection. *Shigella* spp. are Gram-negative bacteria that are the causative agents of shigellosis, a diarrheal disease that disproportionately affects young children in low and middle income countries. *Shigella* causes more than 200 million cases and over 200,000 deaths each year (12). *Shigella* are human-specific pathogens that are transmitted via a fecal-oral route *Shigella* OspF blocks rapid priming of the NAIP–NLRC4 inflammasome and cause disease by invading and replicating within intestinal epithelial cells (IECs). Successful infection requires a plasmid-encoded T3SS which facilitates the cytosolic delivery of ∼30 bacterial effectors into host cells (13).

Although wild-type mice are highly resistant to oral *Shigella* infection, we recently established the first mouse model of infection that uses a physiological oral route of infection (14). This model was established by the genetic elimination of the NAIP–NLRC4 inflammasome, which is highly expressed in mouse intestinal epithelial cells and acts as a potent barrier to *Shigella* infection. Despite the ability of human NAIP–NLRC4 to sense the *Shigella* T3SS needle (MxiH) and rod (MxiI) proteins (4, 5, 14) humans are not protected from infection. One reason for this may be the apparently low expression of NAIP–NLRC4 in human intestinal epithelial cells (15). However, we previously found that the NAIP–NLRC4 inflammasome is suppressed by *Shigella* infection in human THP-1 cells (14). *Shigella* uses many effectors to suppress host immune responses. For example, *Shigella* OspC3 inactivates human CASP4 by ADP-riboxanation (16, 17), and *Shigella* IpaH9.8 degrades anti-bacterial guanylate-binding proteins (18-21). Thus, we hypothesized that *Shigella* might encode an effector that suppresses the NAIP– NLRC4 inflammasome.

In order to test if a secreted *Shigella* effector could suppress NAIP–NLRC4, we conducted a bottom-up screen with a collection of *Shigella* strains that secrete single effectors for those that block NAIP–NLRC4-triggered cell death. This screen identified OspF as a specific suppressor of NAIP–NLRC4. OspF was previously identified as a phosphothreonine lyase that specifically irreversibly inactivates two MAP kinases, p38 and ERK (22-24). We found that p38 rapidly primes NAIP–NLRC4, independent of new transcription and translation, and that OspF inhibits p38-dependent priming of NAIP–NLRC4. These effects were particularly important in cells with low expression of NLRC4. Our results identify a new mechanism of rapid priming of the NAIP–NLRC4 inflammasome, and provide a mechanism by which *Shigella* could evade host immunity provided by NAIP–NLRC4.

## Results

### A mT3Sf screen identifies OspF as a specific suppressor of the NAIP–NLRC4 inflammasome

The T3SS and almost all *Shigella* effectors are encoded on a large 220 kb virulence plasmid (25). mT3Sf is a virulence-plasmid minus variant of *S. flexneri* that contains operons that encode the components needed to form the type III secretion apparatus (T3SA) and four embedded effectors (IpgB1, IpaA, IcsB, IpgD), plus a plasmid that encodes their shared transcriptional regulator VirB under the control of Turcotte et al (2025) *bioRxiv* the P_Tac_ IPTG-inducible promoter. In this background, each mT3Sf strain also harbors a unique plasmid encoding a single effector, or an empty vector (Kim et al, in preparation). These strains express a functional T3SS and are capable of invading and escaping into the cytosol of infected epithelial cells (Kim et al, in preparation). mT3Sf retains the potential to activate the NAIP–NLRC4 inflammasome (via the T3SS rod and needle proteins) and CASP4 (via cytoplasmic LPS) (Figure 1A). Thus, we set out to use the mT3Sf strains to conduct a screen to study the roles of individual effectors in a bottom-up platform where the NAIP–NLRC4 and CASP4 inflammasomes are activated naturally by infection, free of the confounding effects of additional effectors that may normally suppress these inflammasomes or activate others. Infection with mT3Sf::empty induced cell death of infected THP-1 human macrophages, and this death was reduced in *NLRC4/CASP4*^−/−^ double knockout cells to the background levels seen in control *GSDMD*^−/−^ cells (Figure 1B). We observed substantial levels of GSDMD-dependent cell death in mT3Sf-infected *CASP4*^−/−^ cells (mediated by NLRC4) and in *NLRC4*^−/−^ cells (mediated by CASP4), confirming that mT3Sf::empty infection induces pyroptosis via activation of both of these two main cell death pathways. To screen for effector(s) that suppress NAIP–NLRC4, we infected *CASP4*^−/−^ THP-1 cells with a panel of mT3Sf strains, each expressing an individual effector. In these cells, infection with mT3Sf strains expressing OspF, IpaH7.8, IpaH1.4, and OspI significantly suppressed cell death relative to mT3Sf::empty (Figure 1C). To assess whether the inhibition was specific for the NAIP–NLRC4 inflammasome, we also screened the mT3Sf strains in *NLRC4*^−/−^ THP-1 cells (in which cell death is mediated by CASP4). In these cells, we found that OspI, IpaH7.8, IpaH9.8, OspC3, and IpaH1.4 significantly suppressed cell death (Figure 1D).

**Figure 1.**
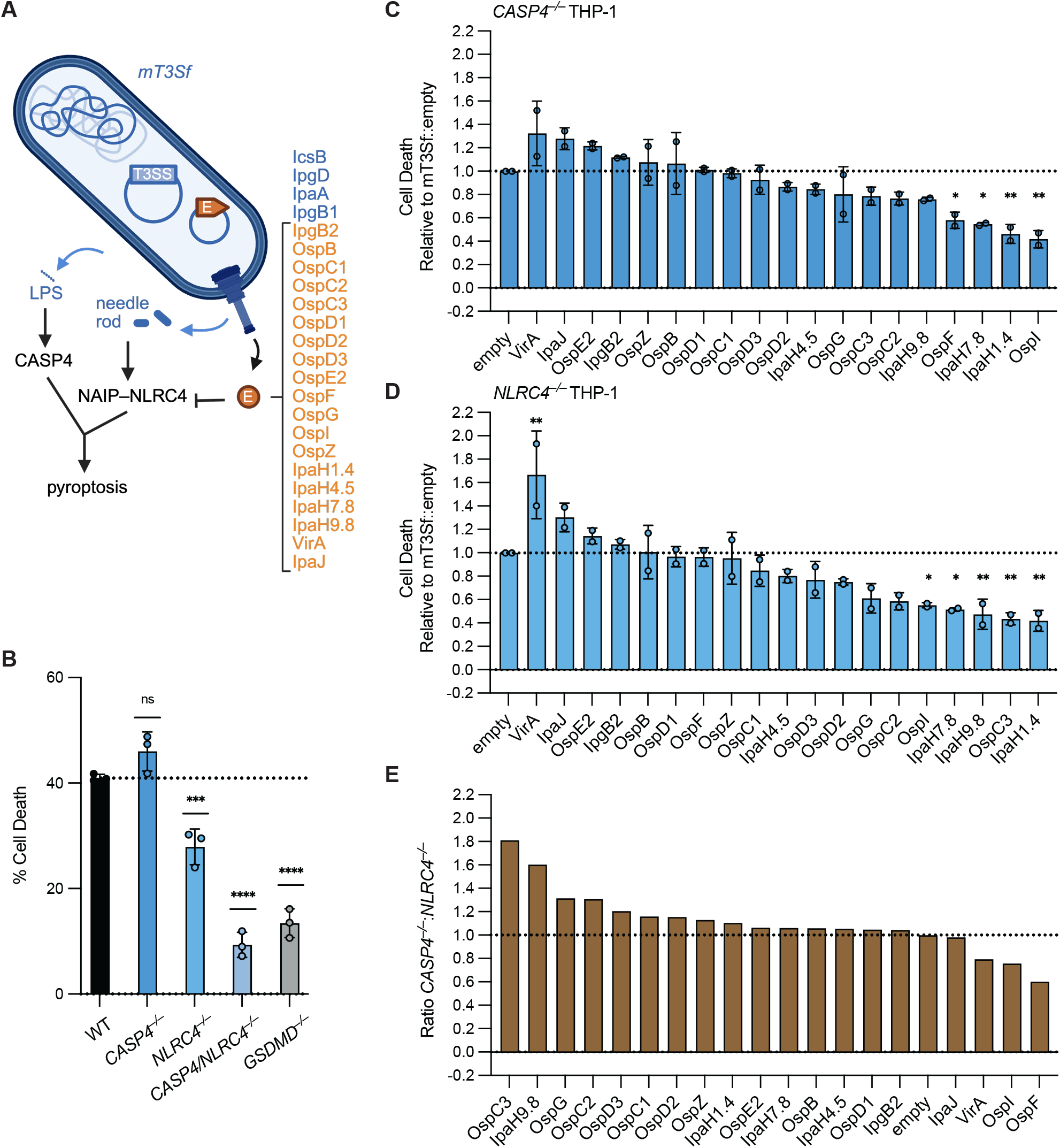
A mT3Sf screen identifies OspF as a specific suppressor of the NAIP–NLRC4 inflammasome. **(A)** mT3Sf model. Features of mT3Sf::empty shown in dark blue with type III secretion system (“T3SS”). Individual effectors (“E”) introduced shown in orange. Created with BioRender.com. **(B)** WT, *CASP4*^−/−^, *NLRC4*^−/−^, *CASP4*/*NLRC4*^−/−^, or *GSDMD*^−/−^ THP-1 cells infected with mT3Sf::empty at an MOI of 5. Cell death was measured at 1 hpi by propidium iodide (PI) uptake and calculated as % Cell Death relative to TritonX-100 treatment. **(C)** *CASP4*^−/−^ or **(D)** *NLRC4*^−/−^ THP-1 cells infected with mT3Sf strains at an MOI of 5 for 2 hr. Percent Cell Death calculated as in B. Data shows cell death relative to mT3Sf::empty infection. **(E)** Ratio of *CASP4*^−/−^:*NLRC4*^−/−^ normalized cell death from C and D. Data shown in B are from one experiment, which are representative of more than three independent experiments. Individual data points represent technical replicates, and shown as mean ± SD. For C and D, data represent the mean ± SD of two experimental repeats of the screen, each point represents the mean of three technical replicates. One-way ANOVA. *P < 0.0332, **P < 0.0021, ***P < 0.0002, ****P < 0.0001.

To identify inhibitors specific for either NLRC4 or CASP4, we calculated the ratio of the cell death between CASP4- and NLRC4-deficient THP-1 cells for each individual mT3Sf::effector strain (Figure 1E). This analysis revealed that some effectors non-specifically suppressed both NLRC4- and CASP4-induced cell death, resulting in a cell death ratio close to 1 (Figure 1E). These effectors include IpaH7.8, which ubiquitylates and induces the degradation of GSDMB and GSDMD (26-28), as well as IpaH1.4 and OspI, which interfere with NF-κ? signaling (29-31). By contrast, OspC3 and IpaH9.8 specifically suppressed CASP4-dependent death in NLRC4-deficient cells, as expected (16-21, 32). Conversely, NLRC4-dependent cell death was specifically and most strongly suppressed by OspF. In addition to suppressing cell death, infection with mT3Sf::OspF also prevented IL-1β processing as compared to mT3Sf::empty (Figure S1A, B). Infection with the mT3Sf::OspF strain also *Shigella* OspF blocks rapid priming of the NAIP–NLRC4 inflammasome suppressed NLRC4-dependent IL-1β processing induced by the synthetic NAIP–NLRC4 agonist NeedleTox. NeedleTox has been previously described (3, 33, 34) and consists of the T3SS needle protein fused to the N-terminal signal sequence of Bacillus anthracis lethal factor (LFn). LFn has no enzymatic or toxic activity, but serves to direct the LFn-Needle fusion protein into the cytosol of host cells via the co-delivered protective antigen (PA) peptide-translocation channel protein. In contrast, OspF did not block the ability of Nigericin to activate NLRP3 (Figure S1A, B). Taken together, we conclude that OspF specifically suppresses NAIP–NLRC4 inflammasome activation during infection.

### OspF suppresses the NAIP–NLRC4 inflammasome during *Shigella* infection via p38 inactivation in human cells

We tested the role of OspF during infection with virulent *Shigella* by infecting THP-1 cells with wild-type (WT), Δ*ospF*, and Δ*ospF*+OspF plasmid complemented *Shigella* strains. Cells infected with the Δ*ospF* mutant showed enhanced cell death compared to WT-infected cells (Figure 2A). Furthermore, this enhanced cell death was entirely NLRC4-dependent (Figure 2A). *CASP4*^−/−^ cells with intact NAIP–NLRC4 responses showed enhanced cell death upon infection with Δ*ospF Shigella*, and complementation with plasmid-encoded OspF fully reversed the enhanced death seen with Δ*ospF* (Figure 2A). OspF is a phosphothreonine lyase that irreversibly removes the phosphorylated hydroxyl moiety from activated mitogen activated protein kinases (MAPKs) by β-elimination (22), with specificity for p38 and ERK1/2 (23, 24). To determine if suppression of NAIP–NLRC4 depends on the same features of OspF that confer its specificity for MAPKs, we complemented Δ*ospF* with a plasmid expressing an OspF KLA mutant, which has mutated lysine and leucine residues to alanine in the N-terminal D-domain that is essential for recognition of MAPK substrates (35). We found that the OspF KLA mutant was unable to reverse the enhanced cell death seen with the Δ*ospF* mutant (Figure 2B). This result indicates that the D-domain is required for OspF suppression of NAIP–NLRC4. We also complemented *Shigella* Δ*ospF* with a plasmid expressing the *Salmonella* OspF homolog, SpvC, another phosphothreonine lyase that targets MAPKs (35, 36). SpvC fully reversed the enhanced cell death produced by infection with the Δ*ospF* mutant (Figure 2B). Thus, inhibition of NAIP–NLRC4 correlates with inhibition of MAPK, and the most parsimonious explanation of our results is that inhibition of NAIP–NLRC4 is via the known ability of OspF to inactivate MAPK. However, it is also possible OspF acts on other substrates.

**Figure 2.**
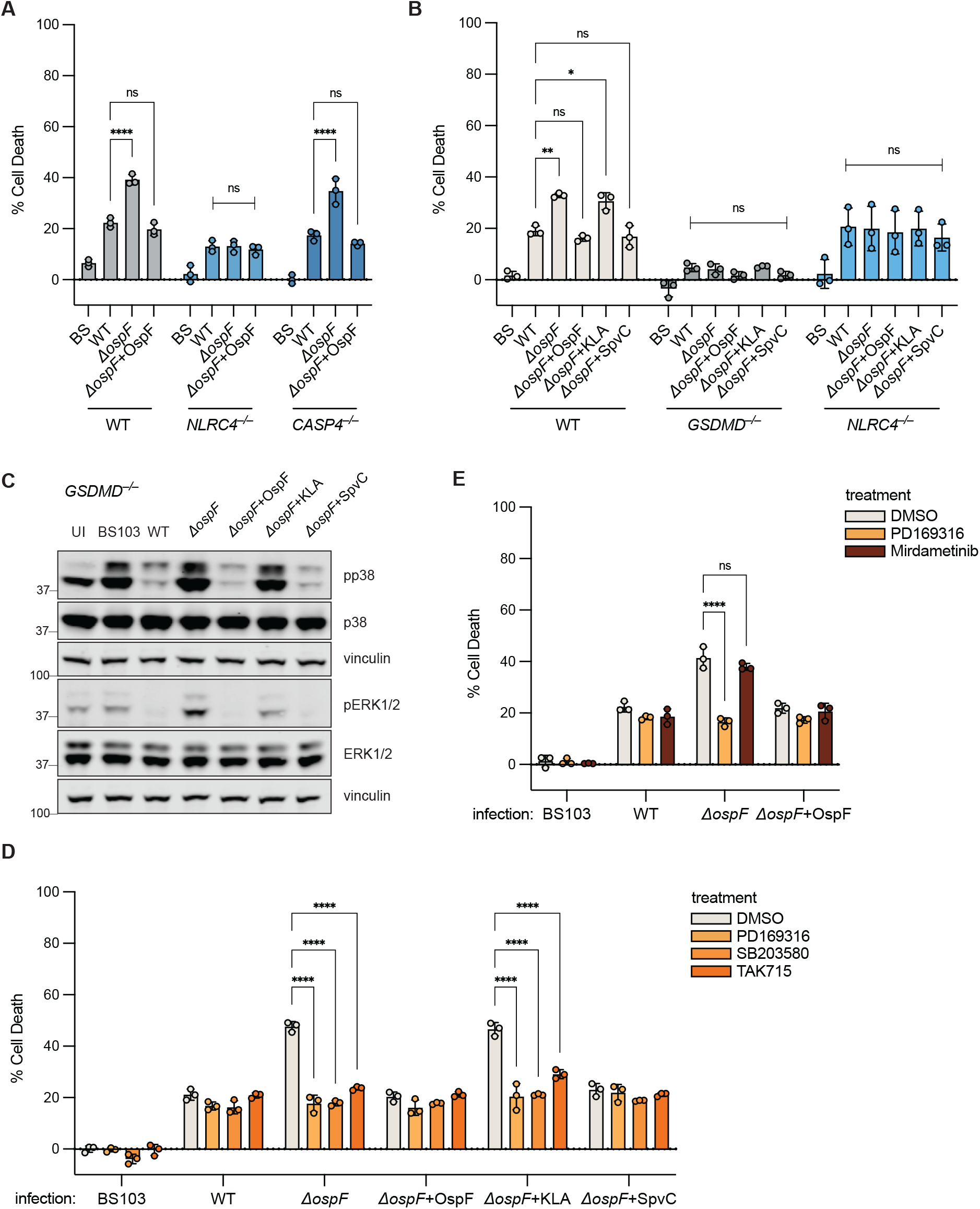
OspF suppresses the NAIP–NLRC4 inflammasome during *Shigella* infection via p38 inactivation in human cells. **(A)** WT, *NLRC4*^−/−^, or *CASP4*^−/−^ THP-1 cells or **(B)** WT, *GSDMD*^−/−^, or *NLRC4*^−/−^ THP-1 cells infected with *Shigella* at an MOI of 10**. C.** Western Blot of lysates from *Shigella*-infected *GSDMD*^−/−^ THP-1 cells, collected at 1 hpi. **(D)** WT THP-1 cells pre-treated for 1 hr with DMSO or p38 inhibitors, PD169316, SB203580, or TAK715 before infection with *Shigella*. **(E)** WT THP-1 cells treated pre-treated for 1 hr with DMSO, PD169316, or Mirdametinib before infection with *Shigella*. Data shown are from one experiment, which are representative of more than three independent experiments. Individual data points represent technical replicates. Cell death was measured at 1 hpi by PI uptake and calculated as % Cell Death relative to TritonX-100 treatment. Data represent the mean ± SD. Two-way ANOVA. *P < 0.0332, **P < 0.0021, ***P < 0.0002, ****P < 0.0001 (A-B, D-E).

To assess if OspF acts via MAPK inhibition, we investigated whether MAPKs are involved in NAIP–NLRC4 activation upon *Shigella* infection. We assayed the induction of phospho-p38 and phospho-ERK1/2 in infected *GSDMD*^−/−^ cells (Figure 2C). In these experiments we used *GSDMD*^−/−^ cells to eliminate the confounding effects of rapid pyroptotic cell death. Consistent with previous work, infection with WT, OspF-complemented, or SpvC-complemented *Shigella* efficiently prevented p38 and ERK1/2 phosphorylation, whereas Δ*ospF*, KLA-complemented, or avirulent (virulence plasmid-cured) BS103 *Shigella* triggered robust MAPK phosphorylation (Figure 2C). To more directly assess the role of MAPK in NAIP–NLRC4 activation, we tested whether MAPK inhibitors blocked *Shigella*-induced NAIP–NLRC4 activation. We found that three different p38 inhibitors (PD169316, SB203580, and TAK715) all prevented NAIP– NLRC4-dependent cell death during Δ*ospF* infection (Figure 2D), whereas use of Mirdametinib to inhibit MEK1/2 upstream of ERK1/2 had no effect (Figure 2E). Furthermore, delivery of *Bacillus anthracis* lethal factor, which cleaves and inactivates MAPK kinases MKK3 and MKK6 upstream of p38 (37-40), eliminated p38 phosphorylation (Figure S2A) and prevented NAIP–NLRC4 activation by Δ*ospF* (Figure S2B). Taken together, these results suggest that OspF suppresses NAIP–NLRC4 indirectly via inactivation of p38 MAPK.

### Rapid priming sensitizes the NAIP–NLRC4 inflammasome in a p38-dependent manner

Our data implied that p38 signaling positively regulates NAIP–NLRC4 inflammasome activation during infection. To test this, we activated p38 using ligands for pattern recognition receptors that are engaged during *Shigella* infection (41-45). Activation of TLR2 by Pam3CSK4, or activation of ALPK1 with ADP-L-Heptose, resulted in robust phospho-p38 in THP-1 cells within 30 minutes (Figure S3A, B). Infection of THP-1 cells with avirulent BS103 also induced a strong phospho-p38 response (Figure S3A). Priming THP-1 cells for just 1 hour before treatment with NeedleTox resulted in a significantly enhanced cell death response compared to unprimed cells (Figure 3A, B). Primed cells responded more quickly than unprimed cells, with significant cell death after just 1 hour of challenge (Figure 3A). Primed cells also responded to a lower dose of NeedleTox to which unprimed cells were unresponsive (Figure 3A, B). As expected, the response of primed cells to NeedleTox was entirely NLRC4-dependent (Figure 3A, B). In addition to increased cell death with priming, we also observed increased conversion of pro-IL-1β to active p17 in rapidly primed *GSDMD*^−/−^ cells in which either the NAIP–NLRC4 or NLRP3 inflammasomes were activated (Figure 3C).

**Figure 3.**
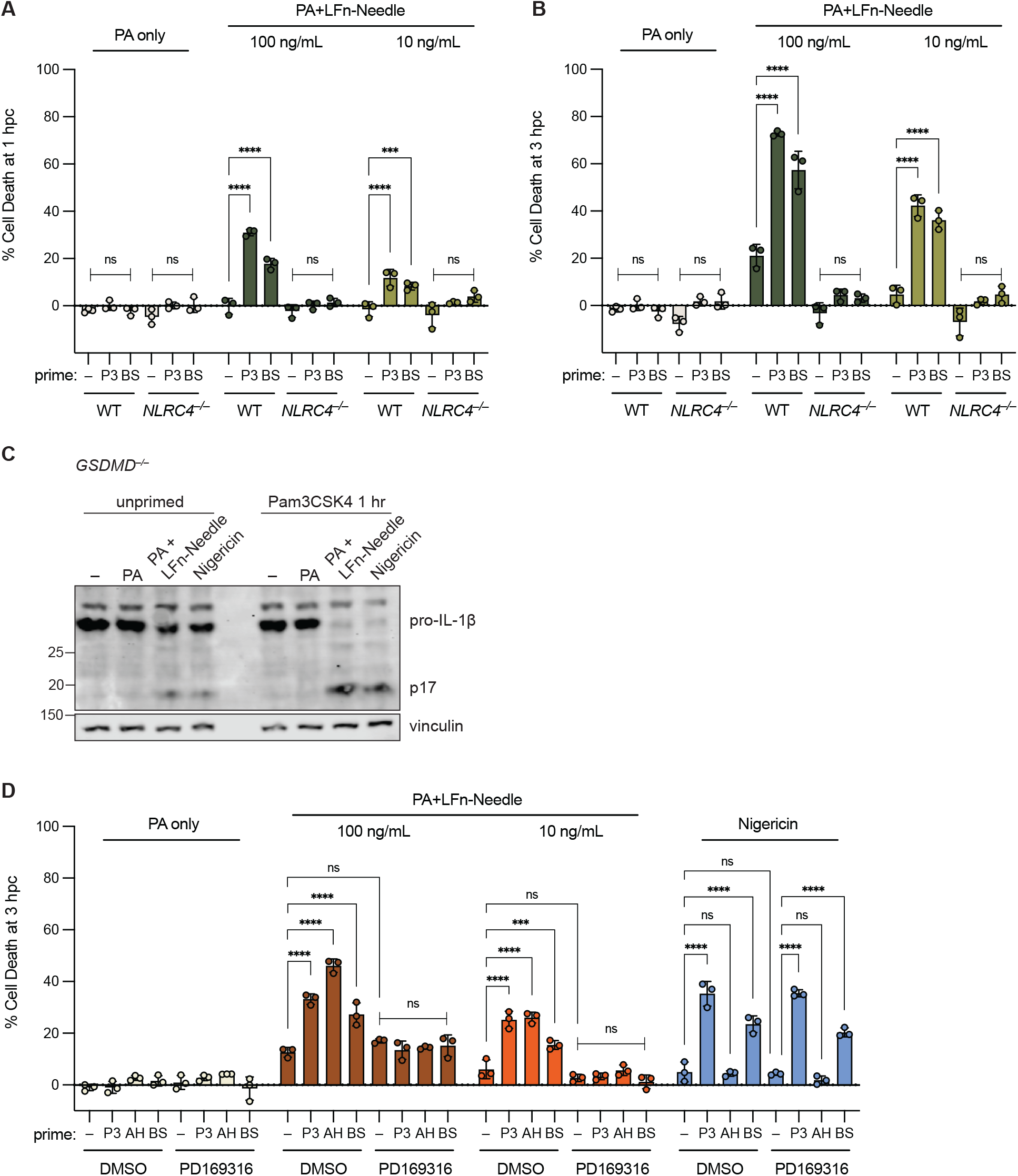
Rapid priming sensitizes the NAIP–NLRC4 inflammasome in a p38-dependent manner. **(A)** and **(B)** WT or *NLRC4*^−/−^ THP-1 cells primed for 1 hr with Pam3CSK4 (P3) or BS103 at MOI 10 (BS) before challenge with PA or PA+LFn-Needle. Cell death collected at 1 hpc (A), and 3 hpc (B). **(C)** Western Blot of lysates from *GSDMD*^−/−^ THP-1 cells primed for 1 hr with P3 or left unprimed before challenge with PA, PA+100 ng/mL LFn-Needle, or Nigericin, collected at 3 hpc. **(D)** WT THP-1 cells pre-treated for 1 hr with DMSO or PD169316, and primed for 1 hr with P3, ADP-L-Heptose (AH), or BS at MOI 10 before challenge with PA, PA+LFn-Needle, or Nigericin. Data shown are from one experiment, which are representative of more than three independent experiments. Individual data points represent technical replicates. Cell death was measured at 1 hpc or 3 hpc by PI uptake and calculated as % Cell Death relative to TritonX-100 treatment. Data represent the mean ± SD. Two-way ANOVA. *P < 0.0332, **P < 0.0021, ***P < 0.0002, ****P < 0.0001.

We found significantly enhanced responses downstream f all ligands that induced a strong phospho-p38 response, including Pam3CSK4, ADP-L-Heptose (Figure 3), and the NOD1 ligand C12-iE-DAP (Figure S3D, E). We also confirmed, as expected, that priming induced by Pam3CSK4 depended on TLR2 and MYD88, whereas priming induced by ADP-L-Heptose depended on ALPK1, and priming induced by C12-iE-DAP depended on NOD1 (Figure S3C-E). Infection with avirulent BS103 *Shigella* primed the NAIP– NLRC4 inflammasome in a TLR2- and MYD88-dependent manner (Figure S3B, C). We cannot test how virulent *Shigella* primes NAIP–NLRC4 since virulent *Shigella* itself induces rapid cell death, but we expect TLR2 ligands to be produced similarly by BS103 and virulent *Shigella*. Furthermore, virulent *Shigella* activates additional sensors, such as NOD1 and ALPK1, which could act redundantly with TLR2 to prime NAIP–NLRC4.

Priming of NAIP–NLRC4 was entirely dependent on p38, as p38 inhibition completely prevented enhanced cell death upon NeedleTox challenge (Figure 3D, Figure S4A). Consistent with the Δ*ospF* infection data, inhibition of ERK1/2 with Mirdametinib had no effect on priming (Figure S4B). Notably, p38 is not required for NAIP–NLRC4 activation, nor does p38 inhibition suppress the NeedleTox responses in unprimed cells (Figure 3D, Figure S4A). Together, these data suggest that p38 is not strictly required for a NAIP–NLRC4 response, but acute p38-dependent priming greatly sensitizes the NAIP–NLRC4 inflammasome to respond.

Notably, although TLR activation also rapidly primed the NLRP3 inflammasome (Figure 3C, D) as previously reported (46-49), ADP-L-Heptose did not rapidly prime NLRP3 (Figure 3D). Nor did p38 inhibitors block rapid NLRP3 priming (Figure 3D). These results indicate that the mechanisms of rapid NLRC4 and NLRP3 priming are distinct. Importantly, priming-enhanced NLRC4-induced cell death was completely blocked by the Caspase-1 inhibitor VX-765 (Figure S5A), indicating that priming promoted canonical Caspase-1 activation downstream of NLRC4 and was not engaging a different Caspase pathway. Furthermore, despite reports that NLRP3 can play a role in NAIP–NLRC4 responses (50), a selective NLRP3 inhibitor MCC950 had no effect on NAIP–NLRC4 priming, indicating that NLRP3 does not contribute to NAIP–NLRC4 priming (Figure S5B).

### Rapid priming enhances NAIP–NLRC4 activation in a transcription- and translation-independent manner and alleviates the requirement for ASC

A recent study showed that extended (16 hours) TLR priming of mouse macrophages induces a p38-dependent up-regulation of NLRC4 expression, resulting in enhanced sensitivity to NAIP ligands (51). However, in this study, human macrophages did not show the same TLR-dependent enhancement of NAIP–NLRC4 responses. In agreement with this prior report, we confirmed that overnight priming with Pam3CSK4 had no effect on NAIP–NLRC4 responses in human THP-1 cells (Figure 4A). However, rapid priming for 1 hour or at the time of challenge significantly enhanced NAIP–NLRC4 responses (Figure 4A). Further examination of short priming incubations showed that Pam3CSK4 incubation for 1 hour or less before challenge had the largest priming effect on NAIP–NLRC4 activity (Figure 4B). After 1 hour, the priming effect became less pronounced and was significantly reduced by 4 hours of Pam3CSK4 treatment, suggesting the priming effect downstream of TLR stimulation is transient in nature. In line with this observation, treatment of cells with the translation inhibitor cycloheximide, or the transcription inhibitor actinomycin D, had no effect on rapid priming of the NAIP–NLRC4 inflammasome (Figure 4C), suggesting priming likely occurs via post-translational modification. Post-translational modifications of inflammasomes have been well-documented (52). One inflammasome component that has been shown to be positively regulated by phosphorylation is the ASC adaptor protein that recruits Caspase-1 downstream of several inflammasomes, including NAIP– NLRC4 (53-56). To determine if rapid priming of NAIP– NLRC4 requires ASC, we challenged *ASC*^−/−^ THP-1 cells under rapid priming conditions. We found that ASC was required for cell death upon NeedleTox challenge in un-primed THP-1 cells. However, rapid priming with Pam3CSK4 alleviated the requirement of ASC for the response to NeedleTox, and primed *ASC*^−/−^ cells were able to undergo NLRC4-induced pyroptosis (Figure 4D). As expected for a PYD-containing NLR, ASC is completely required for NLRP3 inflammasome activation upon nigericin treatment, even with priming (Figure 4D). Thus, ASC is not required for rapid priming of NAIP–NLRC4, and rapid priming allows for ASC-independent inflammasome activation. Phosphorylation of NLRC4 has been described at serine 533. This site has been primarily studied in mice, and its role in NAIP–NLRC4 activation is debated (57-60). In order to test if we could detect phosphorylated S533 (pS533) in our experimental conditions, we used *GSDMD*^−/−^ THP-1 cells overexpressing HA-tagged human or mouse NLRC4 by retroviral transduction with MSCV2.2 constructs. We found that Pam3CSK4 rapid priming was a weak inducer of pS533 in human NLRC4, and that p38 inhibitors were able to prevent the induction of pS533 with Pam3CSK4 priming (Figure S6A, B). Mouse NLRC4 showed stronger induction of phosphorylated species compared to human NLRC4 in all conditions, including unprimed, but as with human NLRC4 phosphorylation was reduced with p38 inhibition (Figure S6A, B).

**Figure 4.**
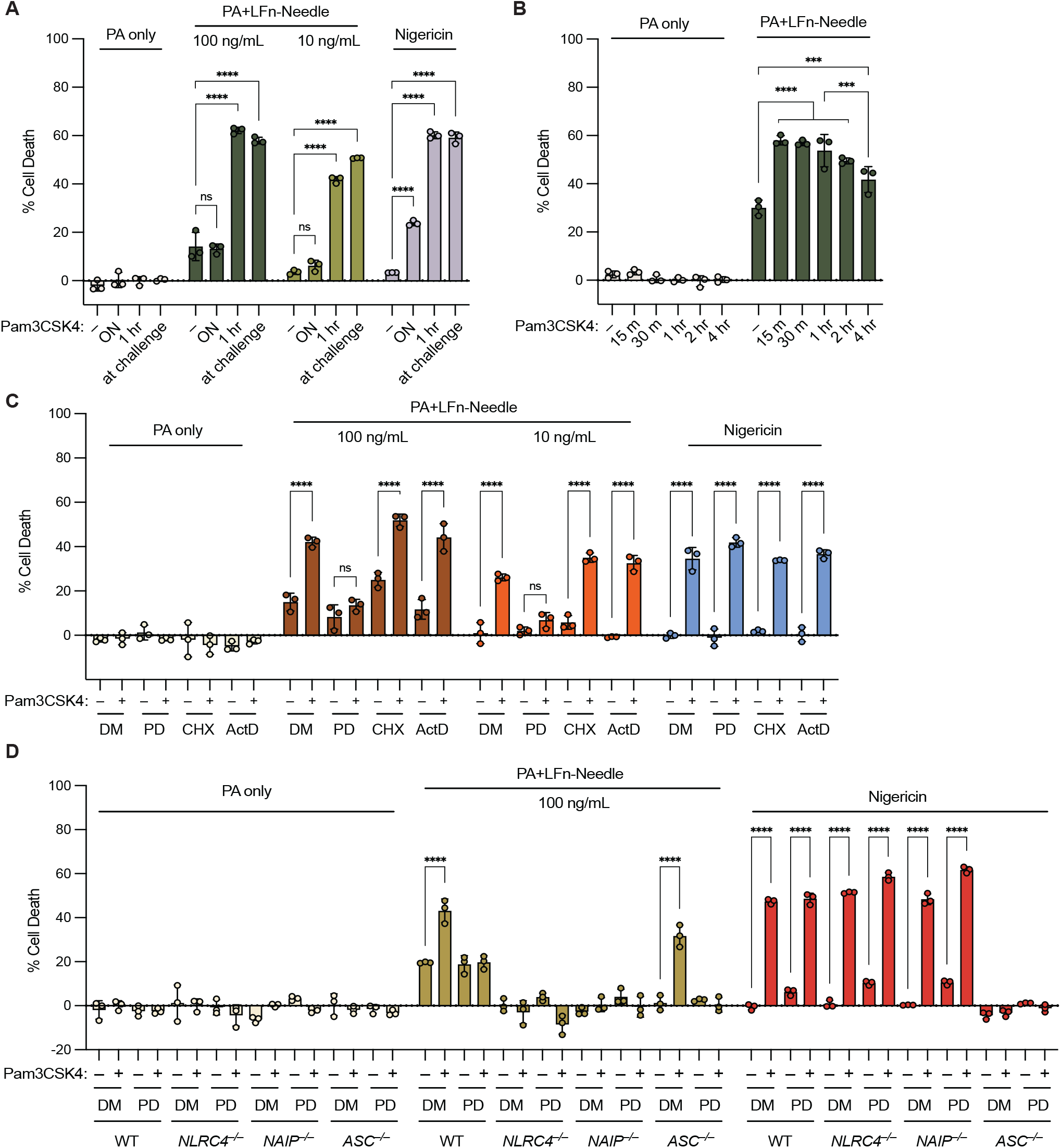
Rapid priming enhances NAIP–NLRC4 activation in a transcription- and translation-independent manner and alleviates requirement for ASC. **(A)** WT THP-1 cells primed with Pam3CSK4 overnight (ON), for 1 hr before challenge, or at the time of challenge. Cells challenged with PA, PA+LFn-Needle, or Nigericin. **(B)** WT THP-1 cells primed with Pam3CSK4 for 15 min (m), 30 m, 1 hr, 2 hr, or 4 hr before challenge with PA or PA+100ng/mL LFn-Needle. **(C)** WT THP-1 cells pre-treated for 1 hr with DMSO (DM), PD169316 (PD), cycloheximide (CHX), or actinomycin D (ActD), and primed for 1 hr with Pam3CSK4 before challenge with PA, PA+LFn-Needle, or Nigericin. **(D)** WT, *NLRC4*^−/−^, *NAIP*^−/−^, or *ASC*^−/−^ THP-1 cells pre-treated for 1 hr with DMSO or PD169316, and primed for 1 hr with Pam3CSK4 before challenge with PA or PA+LFn-Needle, or Nigericin. Data shown are from one experiment, which are representative of more than three independent experiments. Individual data points represent technical replicates. Cell death was measured at 3 hpc by PI uptake and calculated as % Cell Death relative to TritonX-100 treatment. Data represent the mean ± SD. Two-way ANOVA. *P < 0.0332, **P < 0.0021, ***P < 0.0002, ****P < 0.0001.

To study whether phosphorylation of S533 was indeed responsible for the rapid priming effect we generated a NL-RC4_S533A_ knock-in (KI) THP-1 cell line. We found that the NLRC4_S533A_ KI cells lose responsiveness to NeedleTox entirely (regardless of priming status) and phenocopy *NLRC4*^−/−^ THP-1s (Figure S6C), suggesting that the S533A mutation may affect NLRC4 folding or stability. Because we could not use the KI lines to study the role of S533 in priming, we complemented *NLRC4*^−/−^ THP-1 cells with wild-type or S533A NLRC4 using MSCV2.2 retroviral transduction, which results in overexpression of NLRC4. We found that complemented cells were highly sensitized to challenge with NeedleTox, even upon sorting for low-expressing cells (Figure S6D). Despite sensitization to challenge, we could reveal a significant priming effect in the sorted cells challenged with lower doses of NeedleTox. The priming effect was similarly observed in both NLRC4_WT_ and NLRC4_S533A_-complemented cells (Figure S6D). Together these data suggest S533 is not responsible for the observed priming effect regardless of the apparent regulation of this site downstream of p38.

### Rapid priming depends on NLRC4 levels but not NAIP isoform

Our data showed that *NLRC4*^−/−^ THP-1 cells were highly sensitized to NeedleTox challenge when complemented with NLRC4 (Figure 5A, Figure S6D) and this partially masked the priming effect seen in WT THP-1 cells with endogenous levels of NLRC4. Primary human macrophages have been shown to express more NLRC4 and NAIP than THP-1 cells (61). Consistent with this result, we found that primary human monocyte-derived macrophages (HMDMs) are much more responsive to NeedleTox than THP-1 cells, even at 100- and 1000-fold lower doses than typically used to see a response in THP-1 cells (Figure 5B). Despite the high level of responsiveness of primary HMDMs to NeedleTox, we could reveal a modest rapid priming effect with LPS on NAIP–NLRC4 when compiling data from multiple donors (Figure 5B). We sought to examine whether high levels of NLRC4 expression masks the responsiveness of cells to rapid priming. We tested wild-type mouse bone marrow-derived macrophages (BMMs) (Figure 5C), a cell type that is also highly responsive to NLRC4 agonists, and found no evidence of a priming effect. However, when we tested BMMs derived from *Nlrc4*^+/−^ (heterozygous) mice, which express reduced levels of NLRC4, we found that these macrophages were responsive to rapid priming with both Pam3CSK4 and ADP-L-Heptose (Figure 5C). This result suggests that rapid priming is a property of both mouse and human cells, but is apparent only in cells expressing lower NLRC4 levels (THP- 1 or mouse *Nlrc4*^+/−^ BMMs) while being largely or completely masked in primary cells expressing higher levels of NLRC4. To test if human cells expressing mouse NAIPs could be primed, we used retroviral transduction to express mouse *Naip1* and *Naip2* in *NAIP*^−/−^ THP-1 cells (Figure 5D). Mouse NAIP1 detects T3SS Needle and is responsive to NeedleTox (62), whereas mouse NAIP2 detects T3SS rod and is unresponsive to NeedleTox, but is responsive to RodTox (3). THP-1 cells expressing mouse *Naip1* or *Naip2* in place of human NAIP were still regulated by rapid priming (Figure 5D). This result is consistent with the rapid priming we see in mouse *Nlrc4*^+/−^ BMMs and confirm that both human and mouse NAIP proteins can form rapidly primed NAIP– NLRC4 inflammasomes. Furthermore, although overexpression of NLRC4 largely masked the priming effect, cells overexpressing NAIP1 or NAIP2 were still rapidly primed, similar to WT THP-1 cells, suggesting that NLRC4 levels determine the sensitivity to priming.

**Figure 5.**
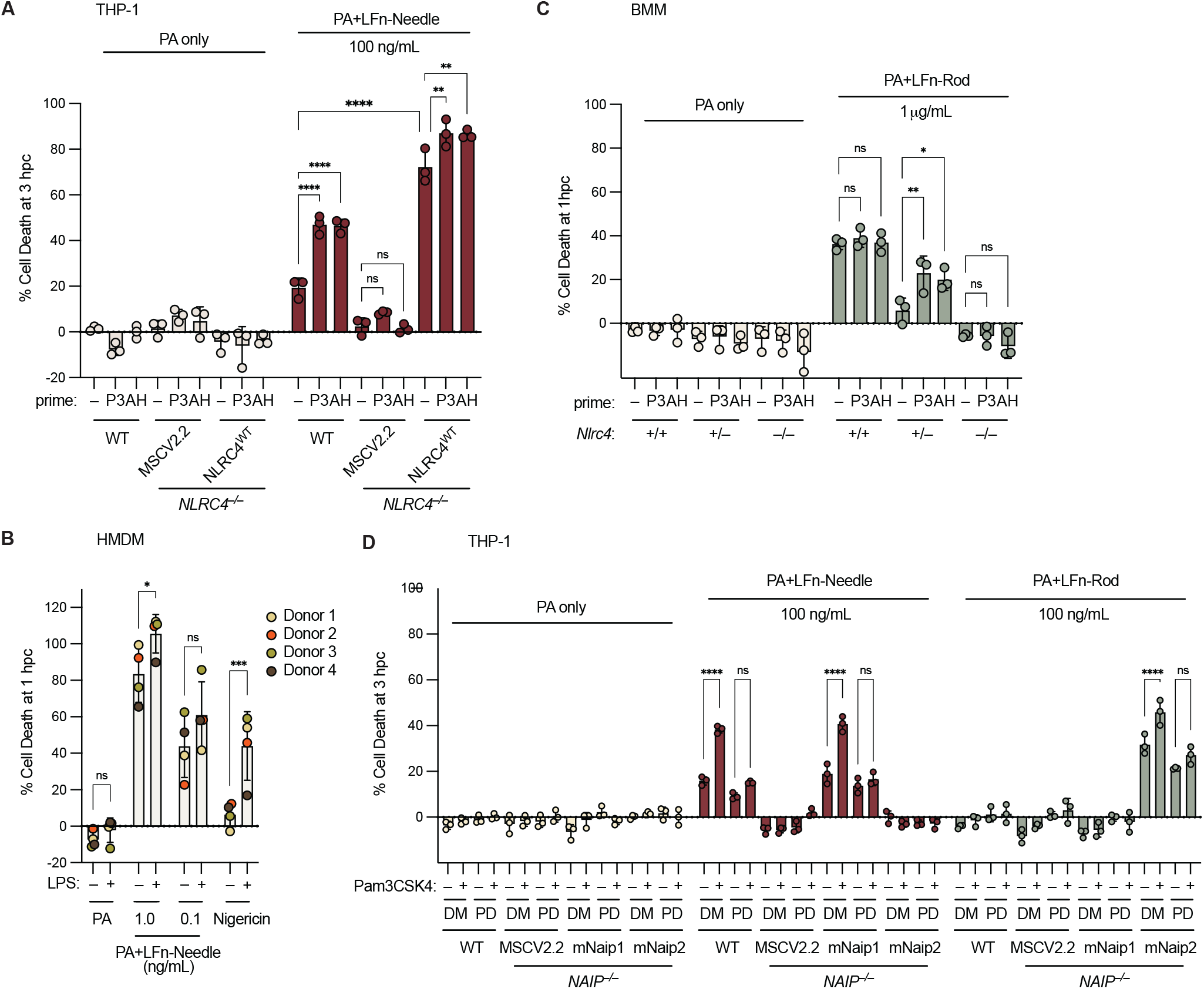
Overexpression of NLRC4, but not NAIP, in THP-1 cells and high expression in primary macrophages masks the priming effect on NAIP–NLRC4. **(A)** WT or transduced *NLRC4*^−/−^ THP-1 cells primed for 1 hr with Pam3CSK4 (P3) or ADP-L-Heptose (AH) before challenge with PA or PA+LFn-Needle **(B)** Primary human monocyte-derived macrophages (HMDM) from four donors primed for 1 hr with LPS before challenge with PA, PA+LFn-Needle, or Nigericin **(C)** Mouse bone marrow-derived macrophages (BMM) primed for 1 hr with P3 or AH before challenge with PA or PA+LFn-Rod **(D)** WT or transduced *NAIP*^−/−^ THP-1 cells pre-treated for 1 hr with DMSO (DM) or PD169316 (PD), and primed for 1 hr with P3 before challenge with PA or PA+LFn-Needle, or PA+LFn-Rod. Data shown are from one experiment, representative of more than three independent experiments. Individual data points represent technical replicates in A, C, and D. In B, each point represents an individual donor (the mean of three technical replicates). Cell death was measured at 1 hpc (B, C) or 3 hpc (A, D) by PI uptake and calculated as % Cell Death relative to TritonX-100 treatment. Data represent the mean ± SD. Two-way ANOVA. *P < 0.0332, **P < 0.0021, ***P < 0.0002, ****P < 0.0001.

## Discussion

Humans are highly susceptible to *Shigella* infection, whereas mice are resistant due to the potent protection afforded by the NAIP–NLRC4 inflammasome. We previously found that NAIP–NLRC4 is actively suppressed during infection of human THP-1 cells (14). Here we found that in human THP-1 cells, NAIP–NLRC4 is suppressed during infection by the secreted effector OspF. Infection with Δ*ospF* resulted in significantly enhanced cell death that depended on both NAIP–NLRC4 and p38 MAP kinases. Our data suggest that the link between p38 and NAIP–NLRC4 activation during *Shigella* infection involves the rapid post-translational priming of NAIP–NLRC4, as treatment of cells for 1 hour or less with innate immune agonists that active p38 resulted in significantly enhanced responses to a NAIP–NLRC4 agonist.

Previous reports identified phosphorylation of serine 533 in NLRC4, although its role in NAIP–NLRC4 activation has remained unclear and less is known about its role in human NLRC4 (57, 58, 60, 63). Previous reports identified PKCδ and LRRK2 as putative kinases regulating pS533 (57, 59, 60), though the role of PKCδ is controversial (64). Our data suggest that S533 is also regulated by or down-stream of p38 MAPK, as p38 inhibitors could prevent phosphorylation at this site under rapid priming conditions (Figure S6A, B). However, our data suggest pS533 is not strictly required for rapid priming, since overexpressed NLRC4_S533A_ could still be primed. Future studies will be important to identify the mechanism of rapid priming. Our attempts to use mass spectrometry to identify additional NLRC4 phosphorylation sites have thus far been inconclusive, and it remains possible that p38 acts indirectly to mediate NLRC4 priming.

Notably, we found that unprimed THP-1 cells required ASC for cell death induced by NeedleTox; however, priming with Pam3CSK4 alleviated the requirement for ASC (Figure 4C). NLRC4 lacks a PYD, but contains a CARD which is capable of direct interaction with the CARD of Caspase-1 (65). Although ASC is generally required for Caspase-1 activation by PYD-containing inflammasomes, the NAIP–NLRC4 inflammasome can induce cell death in the absence of ASC (66-69). Whether ASC is required for NAIP–NLRC4 activation in human cells is not clear. However, it was shown that in the context of *Salmonella* infection in primary human macrophages, ASC is required for cytokine processing but dispensable for pyroptosis, in line with mouse data (70). Consistent with our results, Moghaddas, et al showed that in THP-1 cells, ASC was largely required for both cell death and cytokine response to the T3SS needle protein PrgI (delivered by a retroviral expression vector) (71). In this system, cells were primed with Pam3CSK4 for 3 hours before infection with PrgI-expressing retrovirus, and cell death and cytokine release were measured after 24 hours. Under these conditions, a requirement for ASC was still observed, consistent with our finding that the priming effect on NAIP–NLRC4 is a rapid and transient event. It is possible that the confusion around the role of ASC in both human and murine NAIP–NLRC4 inflammasomes could be explained by the levels of NLRC4 in the cell type used. Primary HMDMs or murine BMMs with high expression, may not require ASC for NAIP–NLRC4-dependent cell death. Low expression in THP-1 cells may dictate the observed ASC-dependence in unprimed conditions. Priming may facilitate assembly, a conformational change in the CARD of NLRC4, or facilitate Caspase-1 recruitment more efficiently. How exactly priming promotes NAIP– NLRC4 activity and ASC-independence will require additional studies.

Another important future area for study is to identify the physiological context in which rapid priming of NAIP– NLRC4 might be important. High NLRC4 expression masks the rapid priming effect, and sensitizes cells to rapid death regardless of priming. Consistent with the NeedleTox challenge experiments, infection of primary HMDMs with *Shigella* undergo robust cell death and show no difference in death between WT and Δ*ospF*-infected cells (Figure S7A), in contrast to what was observed in THP-1 cells. Because primary HMDMs express high levels of NLRC4, p38-dependent rapid priming is not required for a sufficient NAIP– NLRC4 response, thus OspF does not effectively block pyroptosis. Importantly, *Shigella* invades and forms its replicative niche in IECs (not monocytes or macrophages), and in mice, NAIP–NLRC4 in IECs is essential for protection from *Shigella* infection (14, 72). Moreover, NAIP–NLRC4 in mouse IECs has also been shown to be an important contributor for protection against other enteric pathogens (73-75). However, mouse IECs express especially high levels of NLRC4 and NAIPs compared to other cell types, thus likely masking p38-mediated priming of NLRC4 (76). In contrast, it has been reported that NAIP and NLRC4 are expressed at low or undetectable levels in human intestinal organoids or epithelial cell lines (15, 77). However, expression in human epithelial cells within the gut during infection or inflammation may be distinct from those observed in vitro. Indeed, humans with gain-of-function NLRC4 mutations often show gastrointestinal pathology (71, 78-82), suggesting that NLRC4 may function in the gut in vivo in some contexts. Thus, rapid priming may promote inflammasome activation in human epithelial cells. If human IECs express low NAIP–NLRC4 levels, p38-dependent rapid priming would be essential for a protective NAIP–NLRC4 response and would be potently blocked by OspF during infection, as observed in THP-1 cells with low NLRC4 expression. NAIP–NLRC4 function in primary human IECs under rapid priming conditions present an intriguing area for future studies.

Our finding that NAIP–NLRC4 responsiveness can be enhanced by rapid p38-dependent priming adds significantly to our understanding of the regulation of this important inflammasome. The ability of priming to overcome the loss of ASC suggests that priming might be especially critical in the context of a pathogen that inhibits ASC, or in cell types which lack ASC. Moreover, the rapidity and transient nature of the priming suggests that NAIP–NLRC4 may have evolved as a “coincidence detector” in which maximal responsiveness only occurs when the priming (e.g., TLR) signal is coincident with the cytosolic presence of the NAIP ligand (e.g., T3SS Needle). We speculate that such coincidence detection may provide a mechanism to respond preferentially to stimuli—such as infection with a virulent T3SS+ pathogen—that provide both signals within a narrow spatiotemporal window.

## Materials and Methods

### Cell Culture

THP-1 cells were maintained in RPMI with 10% FBS, 100 U/mL penicillin, 100 mg/mL streptomycin, and 2 mM L-glutamine (“complete RPMI”). THP-1 cells were differentiated in 100 ng/mL phorbol myristate acetate (PMA, Invivogen, tlrl-pma) for 48 hr, followed by a 36 hr rest without PMA or antibiotics before use for challenge or infection. THP-1 cells were purchased from ATCC.

GP2-293 cells (Clontech) were grown in DMEM with 10% FBS, 100 U/mL penicillin, 100 mg/mL streptomycin, and 2 mM L-glutamine. Cell lines were tested for mycoplasma by PCR. Primary B6 BMMs were generated by collecting leg bones from WT, *Nlrc4*^+/−^ and *Nlrc4*^−/−^ mice, and crushing the bones by mortar and pestle. Bone marrow was filtered through a 70 µm strainer and treated with ACK lysing buffer (Gibco). Cells were differentiated for 7 days in complete RPMI supplemented with 10% 3T3-MCSF supernatant. BMMs were harvested in cold PBS and replated for experiments. De-identified primary human monocytes were purchased from AllCells. Cryopreserved negatively selected monocytes were thawed and seeded for differentiation in complete RPMI supplemented with 50 ng/mL human M-CSF (PeproTech, 300-25) for 6 days. Cells were collected in trypsin and replated for experiments.

### Bacterial Strains

Experiments were conducted using the WT *S. flexneri* serovar 2a 2457T strain. BS103 is a virulence plasmid-cured strain derived from the WT strain (83). Δ*ospF* and Δ*ospF*+OspF were described previously (24). OspF KLA mutant and *Salmonella typhimurium* SpvC were cloned into pAM238, and are under the control of the OspF promoter via overlap PCR. Complemented strains were grown in presence of spectinomycin. Strains used are listed in Table S1.

### Shigella Infections

Overnight cultures of *Shigella* were grown in 3 mL TSB at 37°C, 220 rpm. On the day of infection *Shigella* cultures were diluted 1:100 in 5 mL TSB and grown for 2-2.5 hr at 37°C until reaching an OD of 0.6-0.8. The bacteria were pelleted and washed with warm complete RPMI without Pen/Strep. Bacteria were added to cells (MOI 10, unless otherwise stated in legend) and spun at 400 xg, for 10 min at 37°C to synchronize infection. The plates were then incubated at 37°C for 10 min before replacing the media with media containing 100 µg/mL propidium iodide (PI) (Sigma) and 100 µg/mL gentamicin (Gibco), and monitored for cell death.

Cultures of mT3Sf were grown and used for infection as with WT *Shigella*, except on the day of infection, mT3Sf were diluted 1:100 in 5 mL TSB and grown for 1 hr. After 1 hr, cultures were supplemented with IPTG (1 mM) and returned to 37°C, 220 rpm until grown for a total of 2.5-3 hr. Infections were performed as described above, and differ only that MOI 5 was used and plates were incubated at 37°C for 20 min before replacing media with PI- and gentamicin-containing media.

### CRISPR-Cas9 Targeting in THP-1s

THP-1 KO lines were generated as described (84). In brief, THP-1 cells were electroporated with a plasmid U6-sgR-NA-CMV-mCherry-T2A-Cas9 plasmid using a Biorad Gene Pulser Xcell. 20 hr after electroporation, mCherry-positive cells were sorted on the BD FACSAria sorter. Sorted cells were plated at limiting dilution to acquire a single cell per well. After 2-3 weeks, clones were selected and submitted for MiSeq. Outknocker analysis was performed and clones with out-of-frame indels were selected for use. Guide sequences were designed using CHOP-CHOP (85). *CASP4*/*NLRC4*^−/−^ and *GSDMD*/*NLRC4*^−/−^ were generated on the *CASP4*^−/−^ and *GSDMD*^−/−^ background, respectively, using the same *NLRC4* gRNA as for the *NLRC4*^−/−^. Guides used are listed in Table S2.

### Retroviral Transduction

GP2-293 cells were reverse-transfected with Lipofectamine 2000 (Invitrogen), VSV-G, and MSCV2.2-IRES-GFP construct. After 18 hr, the media was changed to THP-1 media. 48 hr after media change, the GP2 media was syringe-filtered with a 45 µM filter over THP-1 cells, and 10 µg/mL protamine sulfate was added. Cells were spun at 1000 xg for 1 hr, at 32°C. Transduced cells were expanded and sorted on the BD FACSAria sorter for GFP-positive cells 3-4 days post transduction. MSCV2.2-hNLRC4-HA, MSCV2.2-mNLRC4-HA, MSCV2.2-mNaip1, and MSCV2.2-mNaip2 have been described previously (3, 14). MSCV2.2-hNLRC4S533A-HA was generated using Q5 Site-Directed Mutagenesis (New England Biolabs).

### Cell Treatment and Inhibitor Conditions

For activating PRRs, cells were treated with 100 ng/mL Pam3CSK4 (Invivogen, tlrl-pms), 100 ng/mL ADP-L-Hep-tose (Invivogen, tlrl-adph-l), 100 ng/mL LPS (Enzo, ALX-581-013), or 1 µg/mL C12-iE-DAP (Invivogen, tlrl-c12dap) for the indicated times. Inhibitors were added 1 hr before infection or challenge and were present throughout the experiment. For NLRP3 activation, cells were treated with 5 µg/mL Nigericin (Invivogen, tlrl-nig). Cells were treated with 5 µM PD169316 (MCE, HY-10578), 5 µM SB203580 (SelleckChem, S2928), 5 µM TAK715 (SelleckChem, S1076), 5 µM Mirdametinib (MCE, HY-10254), 20 µM VX-765 (Invivogen, inh-vs765i-1), 10 µM MCC950 (Invivogen, inh-mcc), 250 ng/mL cycloheximide (Sigma), or 2 µg/mL Actinomycin D (ThermoFisher).

### PA Delivery

Anthrax Lethal Factor protease is delivered into cells via co-administration with protective antigen (PA). Cells were treated with 2 µg/mL PA (List Labs, 171E) only as a control, or 2 µg/mL PA + 1 µg/mL Lethal Factor (List Labs, 169L). Ligands fused to the N-terminal domain of anthrax Lethal Factor (LFn) can be translocated into the cytosol only when administered with PA (3, 33, 34). For NAIP–NLRC4 activation, the combination of both PA and LFn-Needle or LFn-Rod (known as NeedleTox or RodTox) allows for cytosolic delivery of NAIP–NLRC4 ligands and activation. PA, LFn-Needle, or LFn-Rod alone do not induce cell death. Cells were treated with 2 µg/mL PA only, or 2 µg/mL PA + LFn-Needle (Invivogen, tlrl-ndl), or PA + LFn-Rod (Invivogen, tlrl-rod) at the indicated concentrations of LFn proteins.

### Cell Death Assay

For cell death assays, cells were plated in 96 well format (in triplicate) at 50,000 cells per well. To measure cell death, 100 µg/mL PI (Sigma) was added at the time of challenge with inflammasome activators, or at gentamicin addition for infection experiments. PI uptake was measured using a Tecan Spark plate reader. Cell death was calculated relative to 1% Triton X-100-treated wells (100% cell death), and all subtracted background from media only controls.

### Western Blot

Whole-cell lysates were prepared by lysis in RIPA buffer (1% NP-40, 0.1% SDS, 0.5% sodium deoxycholate, 50 mM Tris, 150 mM NaCl, 5 mM EDTA) with freshly supplemented HALT protease and phosphatase inhibitor cocktail (ThermoFisher) for 20 min on ice. Lysates were cleared by centrifugation at 18,000 xg for 20 min at 4°C.

Laemmli buffer was added and boiled for 10 min at 95°C. Samples were separated on NuPAGE Bis-Tris 4-12% gels (ThermoFisher) and transferred onto Immobilon-FL PVDF membranes. Membranes were blocked with Li-Cor Intercept blocking buffer. Primary antibodies were incubated overnight at 4°C. Secondary Li-Cor IRDye-conjugated antibodies were used at 1:5000. Blots were imaged using Li-Cor Odessey CLx. For p38 blots, after probing for phospho-p38, blots were stripped with Li-Cor NewBlot IR stripping buffer before probing with anti-p38 antibody.

Primary antibodies used are listed in Table S3.

### Statistical Analysis

Data were analyzed using GraphPad Prism 10. Statistical tests are indicated in the figure legends.

## Acknowledgments

We would like to thank the Vance and Barton lab members for helpful discussions, and Dr. Sunny Shin (University of Pennsylvania) for an NLRC4 construct. This study was funded by National Science Foundation Graduate Research Fellowship Program DGE 1752814 (E.A.T.). R.E.V. is supported by an Investigator Award and an Emerging Pathogens Initiative Award from the Howard Hughes Medical Institute. R.E.V. is also supported by NIH grants AI075039 and AI155634. C.F.L. acknowledges research support from NIH grant AI169795.

## Author contributions

E.A.T, P.S.M., and R.E.V. conceptualized the project.

E.A.T. performed the experiments and analyzed the results. K.K., L.G., and C.F.L. established and provided mT3Sf strains, and provided *Shigella* strains. K.D.E. provided mice. C.F.L. and R.E.V. advised the project. E.A.T. and R.E.V. wrote the manuscript with feedback from all authors.

## Competing interests

R.E.V. is a scientific advisor to X-biotix Therapeutics, Remedy Plan, and Ditto Biosciences.

## Supplemental Figures and Tables

**Figure S1.**
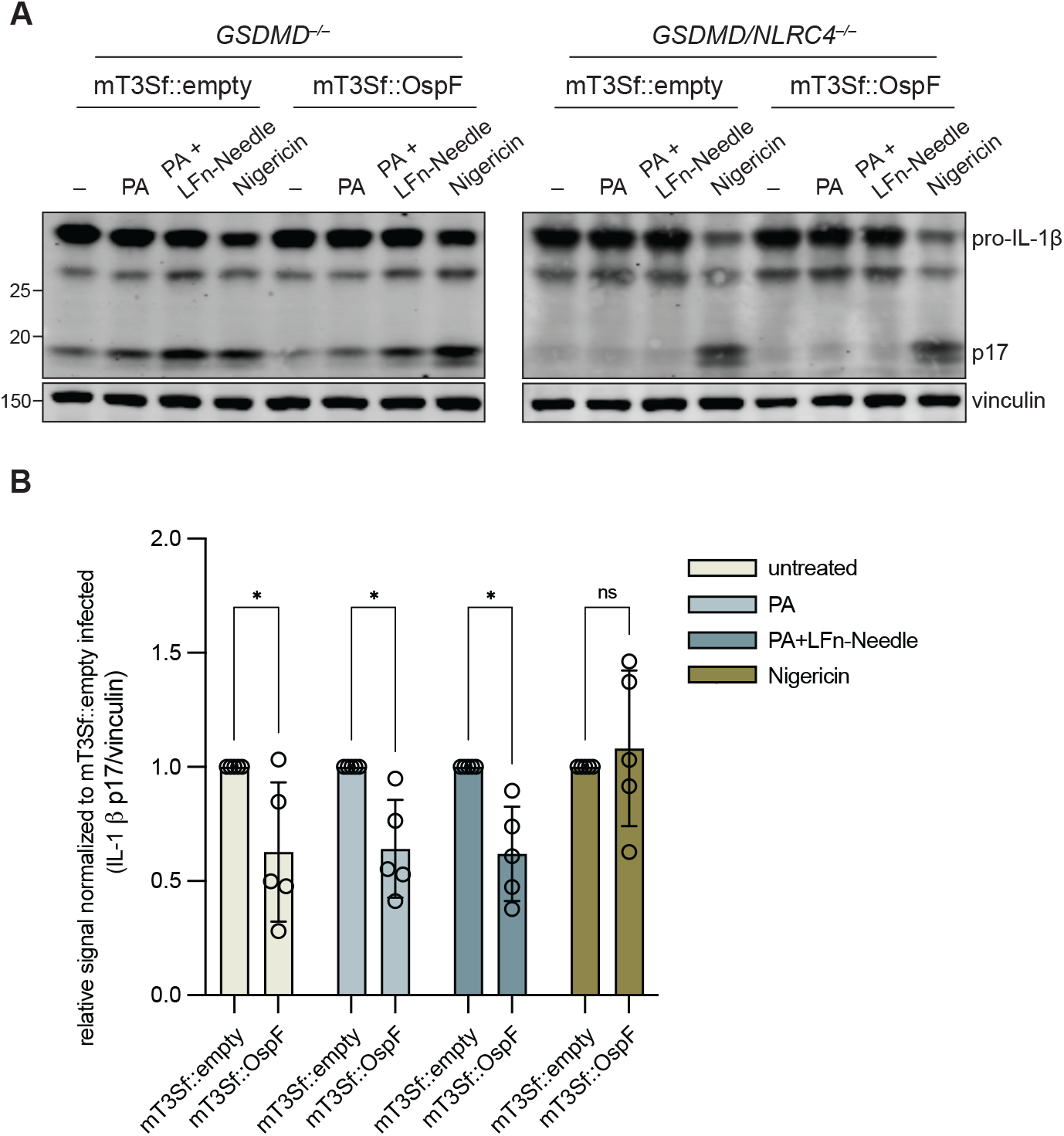
mT3Sf::OspF infection suppresses NLRC4-dependent IL-1β processing. Western Blot of lysates from *GSDMD*^−/−^ or *GSDMD/NLRC4*^−/–^ (**A**) and quantification (**B**) of A of *GSDMD*^−/–^ THP-1 cells infected at MOI 5 with mT3Sf::empty or mT3Sf::OspF for 1 hr before challenge with PA, PA + 100 ng/mL LFn-Needle, Nigericin, or left unchallenged for 2 hr. Data are representative of five independent experiments. Quantification represents p17 band signal normalized by vinculin, and shown relative to mT3Sf::empty-infected band for each challenge condition. Data represent the mean ± SD. Two-way ANOVA. *\*P < 0.0332, **P < 0.0021, ***P < 0.0002, ****P < 0.0001.*

**Figure S2.**
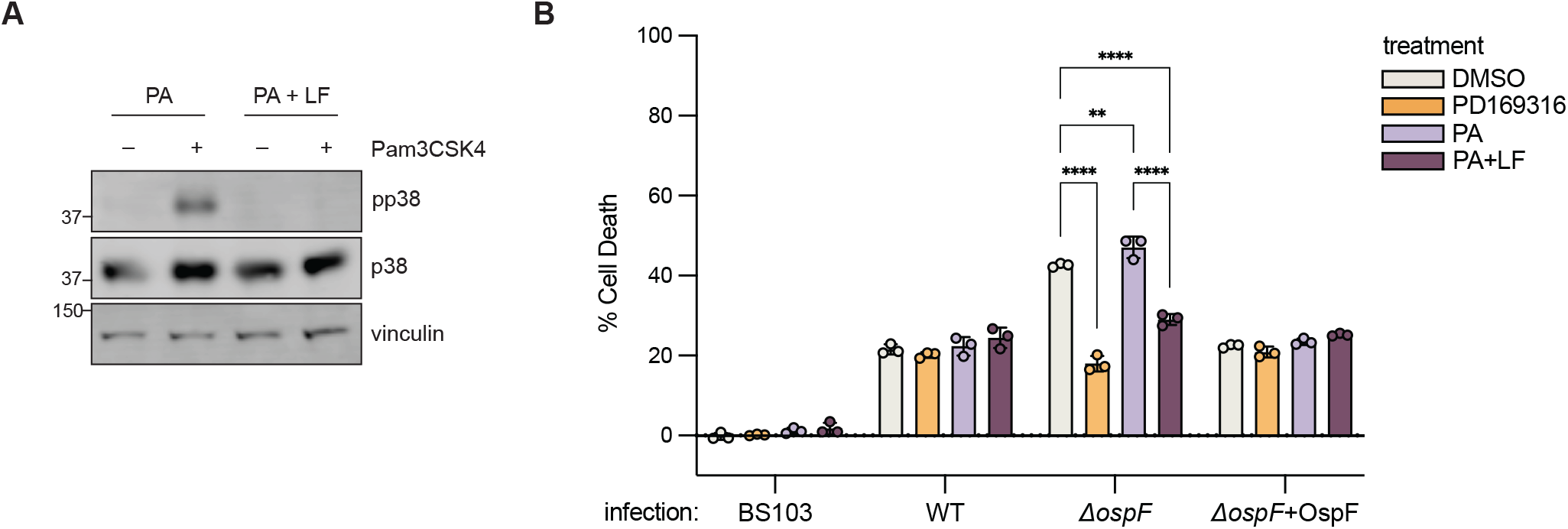
Lethal Factor inactivation of MKKs suppresses NLRC4-dependent cell death during *ΔospF* infection. **A.** Western Blot of lysates from WT THP-1 cells treated with PA or PA + Lethal Factor (LF) for 1 hr before priming with Pam3CSK4 for 1 hr. **B.** WT THP-1 cells treated pre-treated for 1 hr with DMSO, PD169316, PA, or PA + LF before infection with *Shigella* at an MOI of 10. Data shown are from one experiment, which are representative of more than three independent experiments. Individual data points represent technical replicates. Cell death was measured at 1 hpi by PI uptake and calculated as % Cell Death relative to TritonX-100 treatment. Data represent the mean ± SD. Two-way ANOVA. *\*P < 0.0332, **P < 0.0021, ***P < 0.0002, ****P < 0.0001* (A-B, D-E).

**Figure S3.**
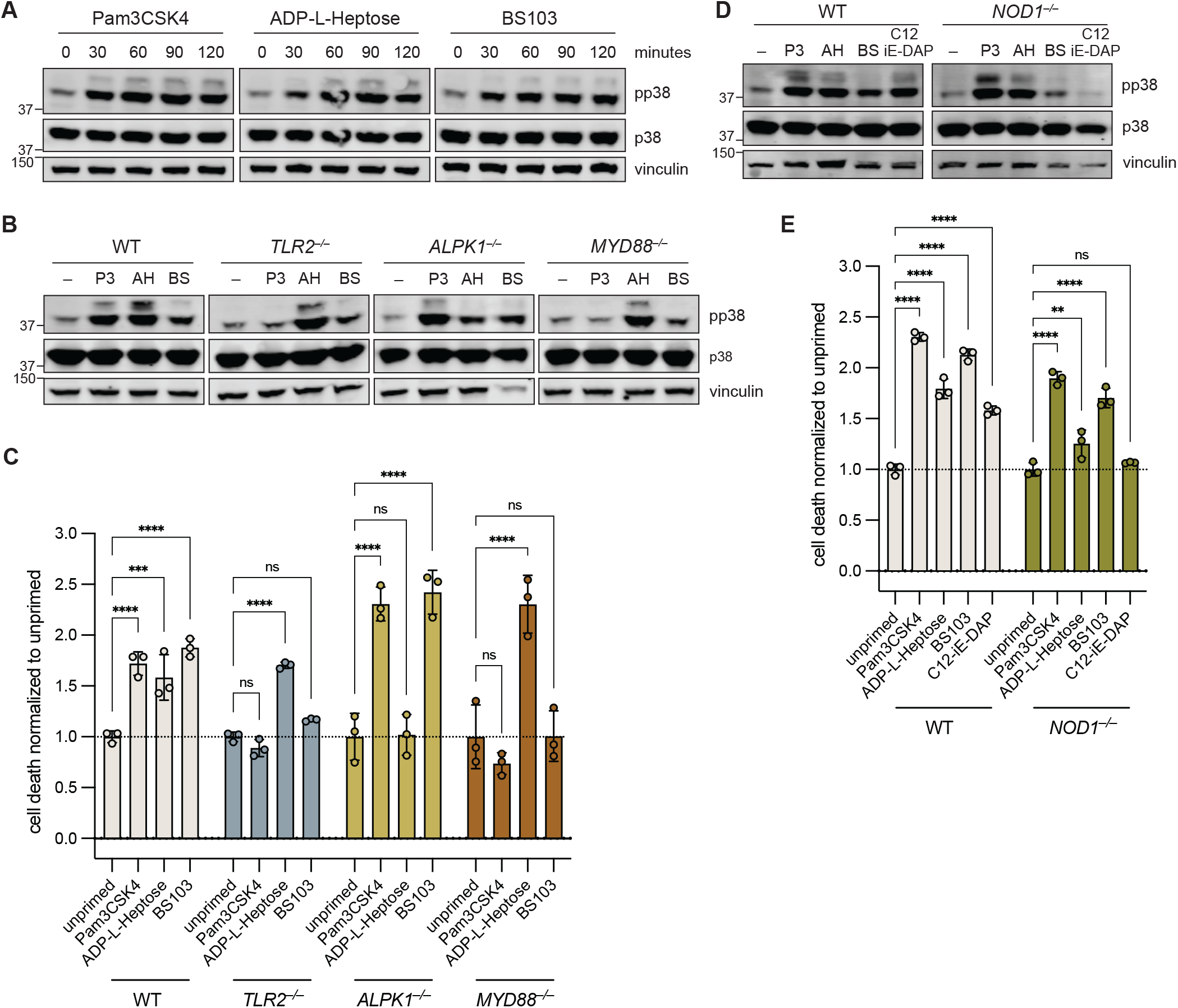
TLR2, ALPK1, and NOD1 activation can prime NAIP–NLRC4. **A.** Western Blot of lysates from priming timecourse of WT THP-1 treated with Pam3CSK4, ADP-L-Heptose, or BS103. **B.** and **C.** THP-1 knockout validation of *TLR2, ALPK1*, and *MYD88^−/−^* THP-1 cells for priming induced phospho-p38 (pp38) (B), and enhanced response to NeedleTox challenge (C). **D.** and **E.** THP-1 knockout validation of *NOD1^−/−^* THP-1s for priming induced pp38 (D), and enhanced response to NeedleTox challenge (E). C and E Cell death was measured at 3 hpc by PI uptake and calculated as % Cell Death relative to TritonX-100 treatment. Fold change of cell death of primed cells relative to unprimed cells, all treated with NeedleTox. Data shown are from one experiment, which are representative of more than three independent experiments. Individual data points represent technical replicates. Data represent the mean ± SD. Two-way ANOVA. *\*P < 0.0332, **P < 0.0021, ***P < 0.0002, ****P < 0.0001.*

**Figure S4.**
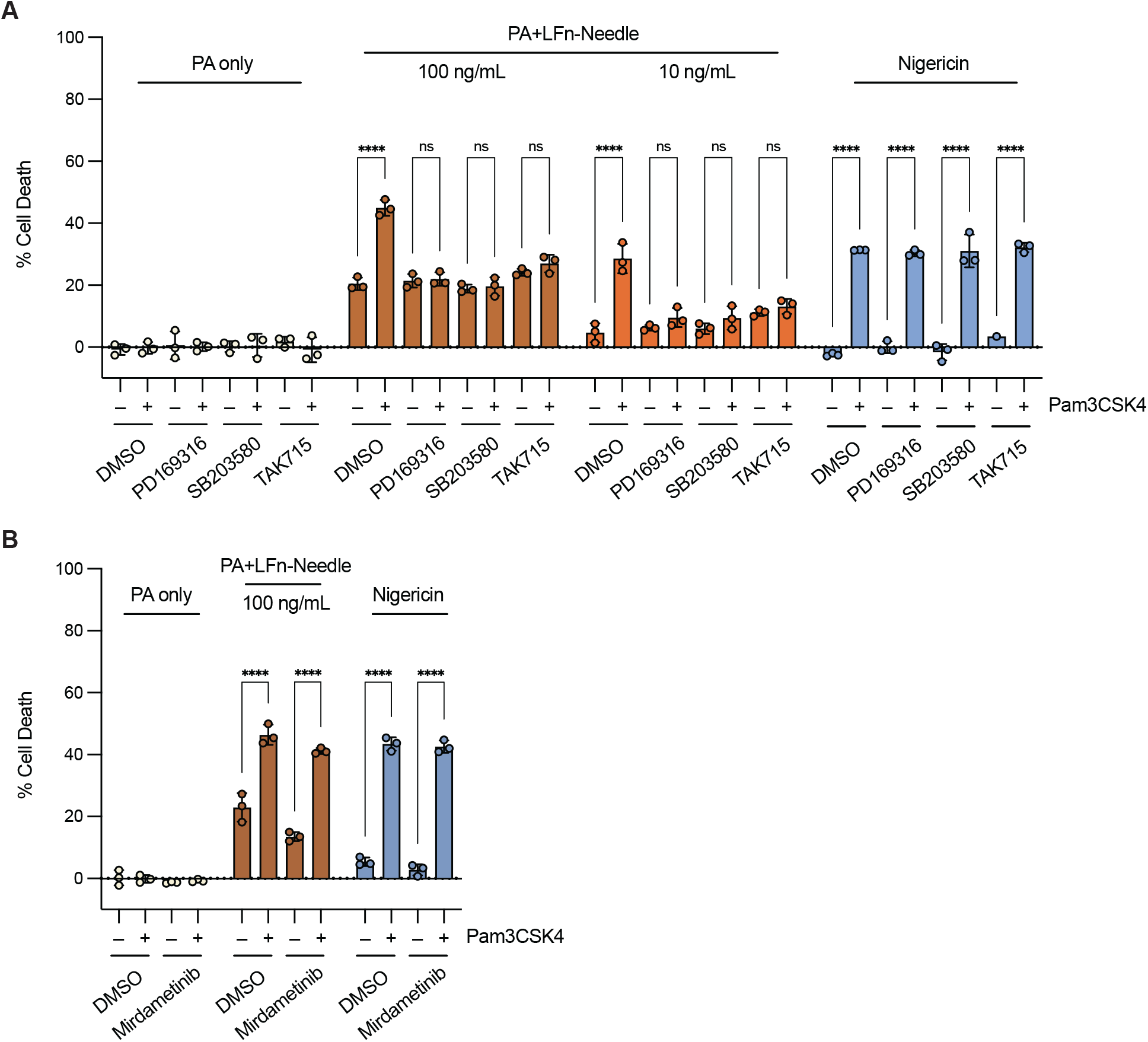
p38 inhibitors suppress rapid priming of NAIP–NLRC4. **A.** WT THP-1 cells pre-treated for 1 hr with DMSO or p38 inhibitors, PD169316, SB203580, or TAK715 before challenge with PA, PA+LFn-Needle, or Nigericin. **B.** WT THP-1 cells pre-treated for 1 hr with DMSO or MEK1/2 inhibitor mirdametinib before challenge with PA, PA+LFn-Needle, or Nigericin. Data shown are from one experiment, which are representative of more than three independent experiments. Individual data points represent technical replicates. Cell death was measured at 3 hpc by PI uptake and calculated as % Cell Death relative to TritonX-100 treatment. Data represent the mean ± SD. Two-way ANOVA. *\*P < 0.0332, **P < 0.0021, ***P < 0.0002, ****P < 0.0001.*

**Figure S5.**
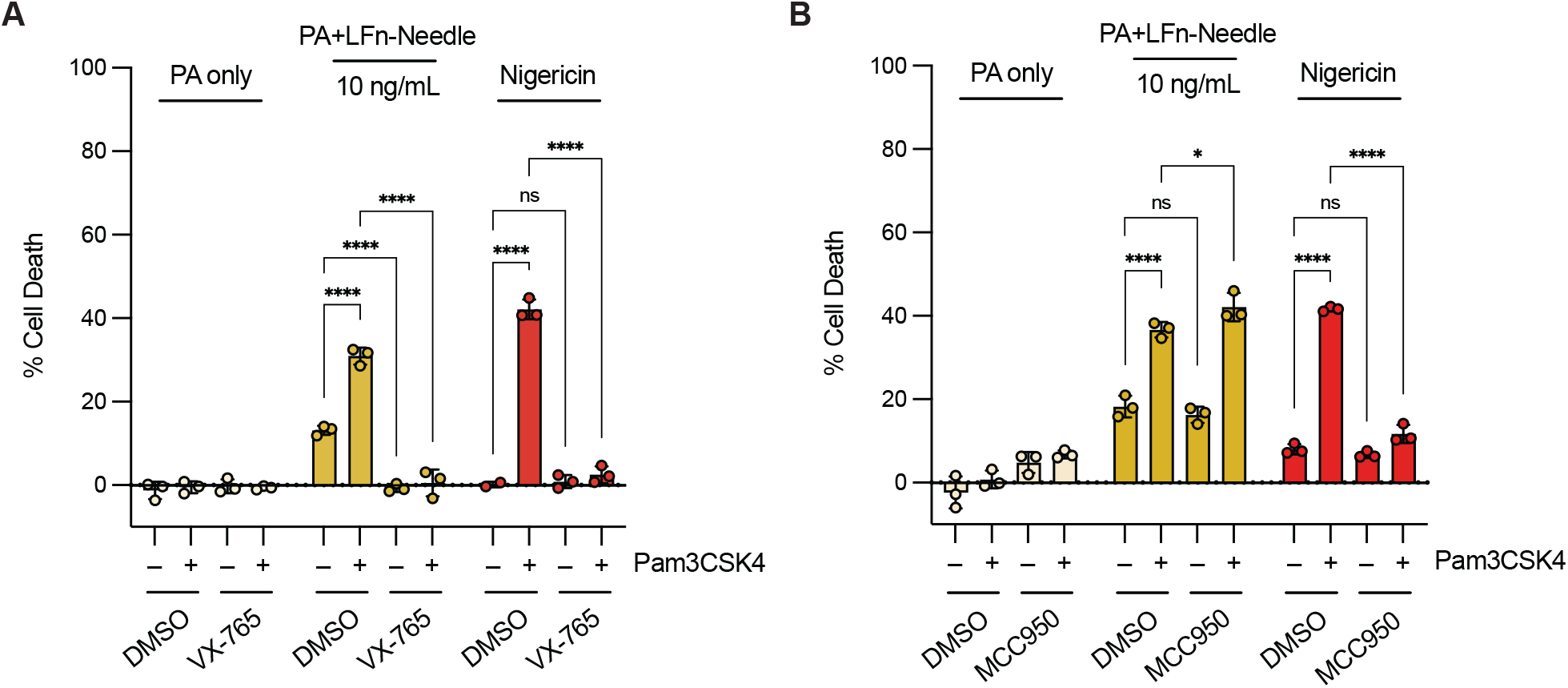
Priming of NAIP–NLRC4 is CASP1-dependent and NLRP3-independent. **A.** WT THP-1 cells pre-treated for 1 hr with DMSO or 20 µM Caspase-1 inhibitor VX-765 before challenge with PA, PA+LFn-Needle, or Nigericin. **B.** WT THP-1 cells pre-treated for 1 hr with DMSO or 10 µM NLRP3 inhibitor MCC950 before Pam3CSK4 prime for 1 hr and challenge with PA, PA+LFn-Needle, or Nigericin. Data shown are from one experiment, which are representative of more than three independent experiments. Individual data points represent technical replicates. Cell death was measured at 3 hpi by PI uptake and calculated as % Cell Death relative to TritonX-100 treatment. Data represent the mean ± SD. Two-way ANOVA. *\*P < 0.0332, **P < 0.0021, ***P < 0.0002, ****P < 0.0001* (A).

**Figure S6.**
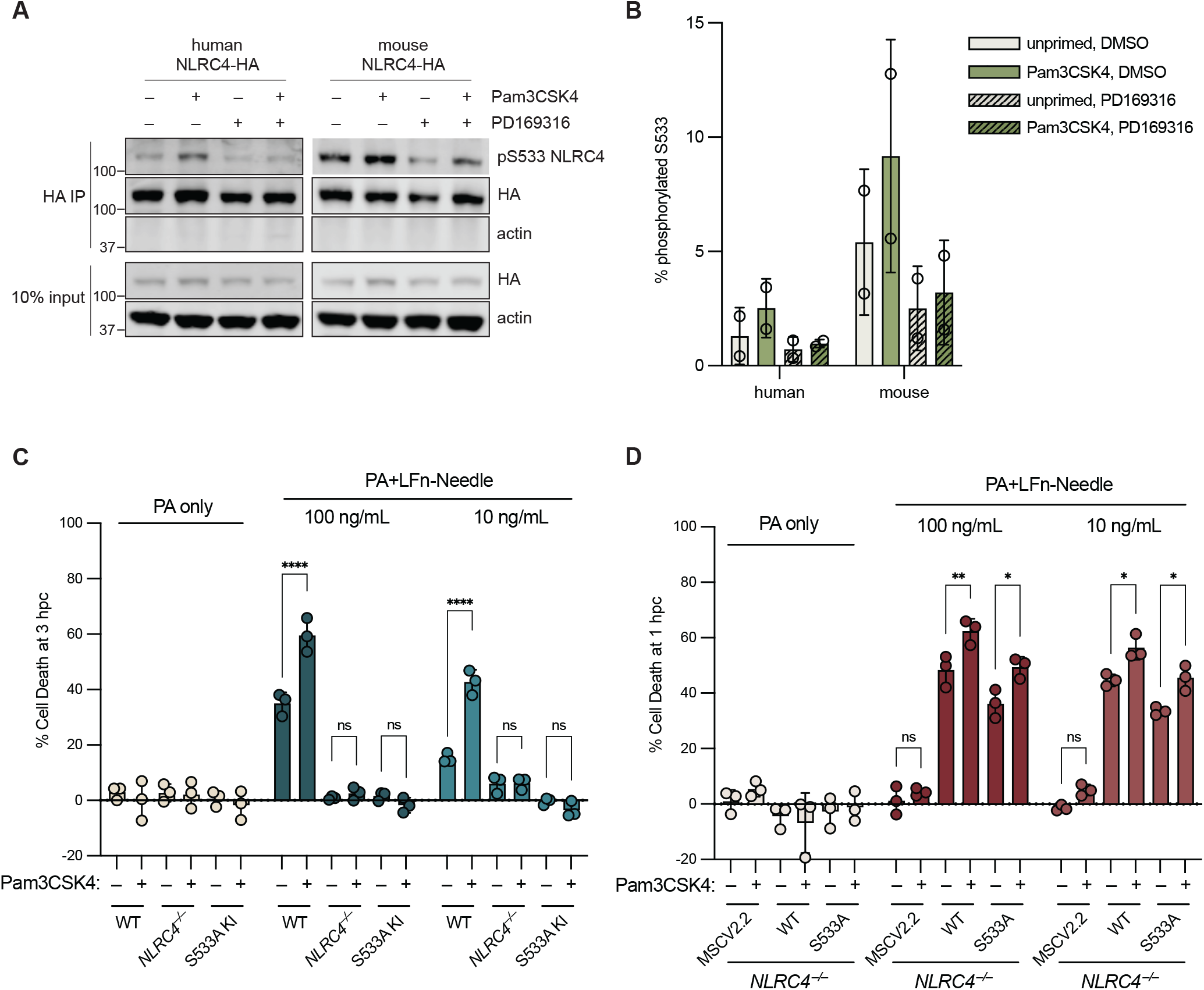
Phosphorylation of NLRC4 S533 is regulated by p38 but is not required for rapid priming. **A.** Western Blot of HA IP of NLRC4-HA in *GSDMD*^−/−^ THP-1 cells upon DMSO or 5 μM PD169316 treatment for 1 hr before prime with Pam3CSK4 for 1 hr. **B.** Quantification of % pS533 NLRC4. Phosphorylated S533 band/total HA-tagged NLRC4. **C.** WT, *NLRC4^−/−^*, or S533A knock-in (KI) THP-1 cells were primed for 1 hr with Pam3CSK4 before challenge with PA only or NeedleTox. **D.** Sorted THP-1 cells with low expression of MSCV2.2 empty or MSCV2.2 NLRC4^WT^ or NLRC4^S533A^ were primed for 1 hr with Pam3CSK4 before challenge with PA only or NeedleTox. Data shown are from one experiment, which are representative of more than three independent experiments. Individual data points represent technical replicates (C and D). Cell death was measured at 3 hpc (**C**) or 1 hpc (**D**) by PI uptake and calculated as % Cell Death relative to TritonX-100 treatment. Data represent the mean ± SD. Two-way ANOVA. *\*P < 0.0332, **P < 0.0021, ***P < 0.0002, ****P < 0.0001.*

**Figure S7.**
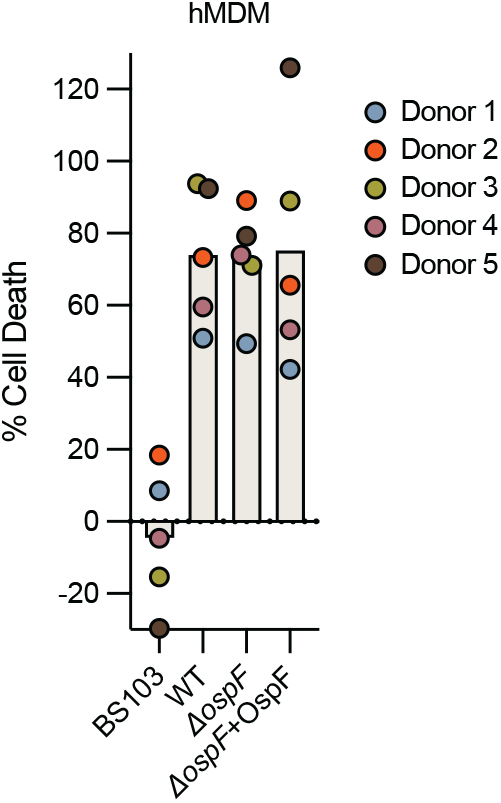
Cell death during primary HMDM infection is not suppressed by OspF. Primary HMDMs infected with *Shigella flexneri* (MOI 5). Each point represents an individual donor (the mean of three technical replicates). Cell death was measured at 1 hpi by PI uptake and calculated as % Cell Death relative to TritonX-100 treatment.

**Table S1.**
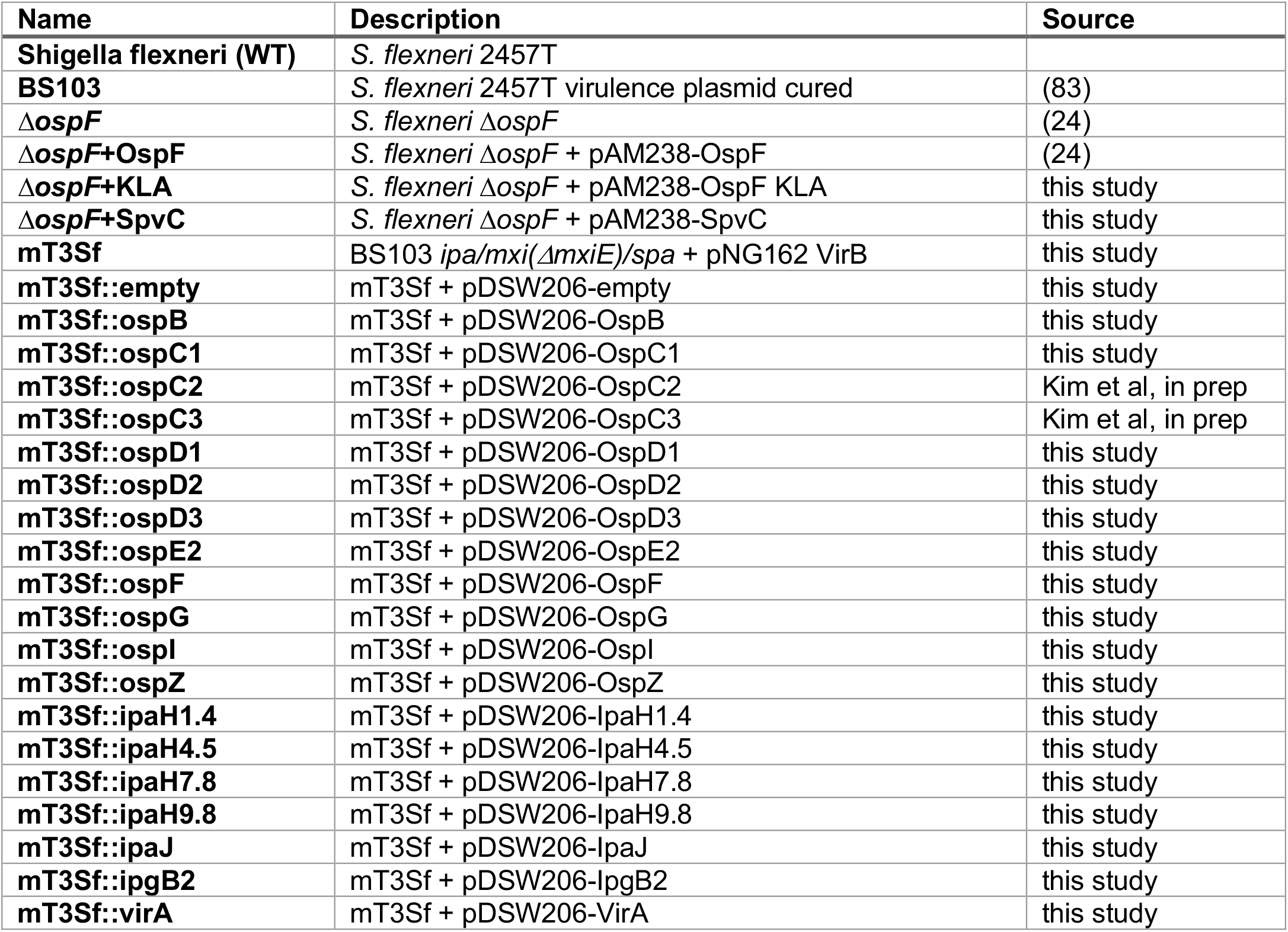
Bacterial strains used in this study.

**Table S2.**
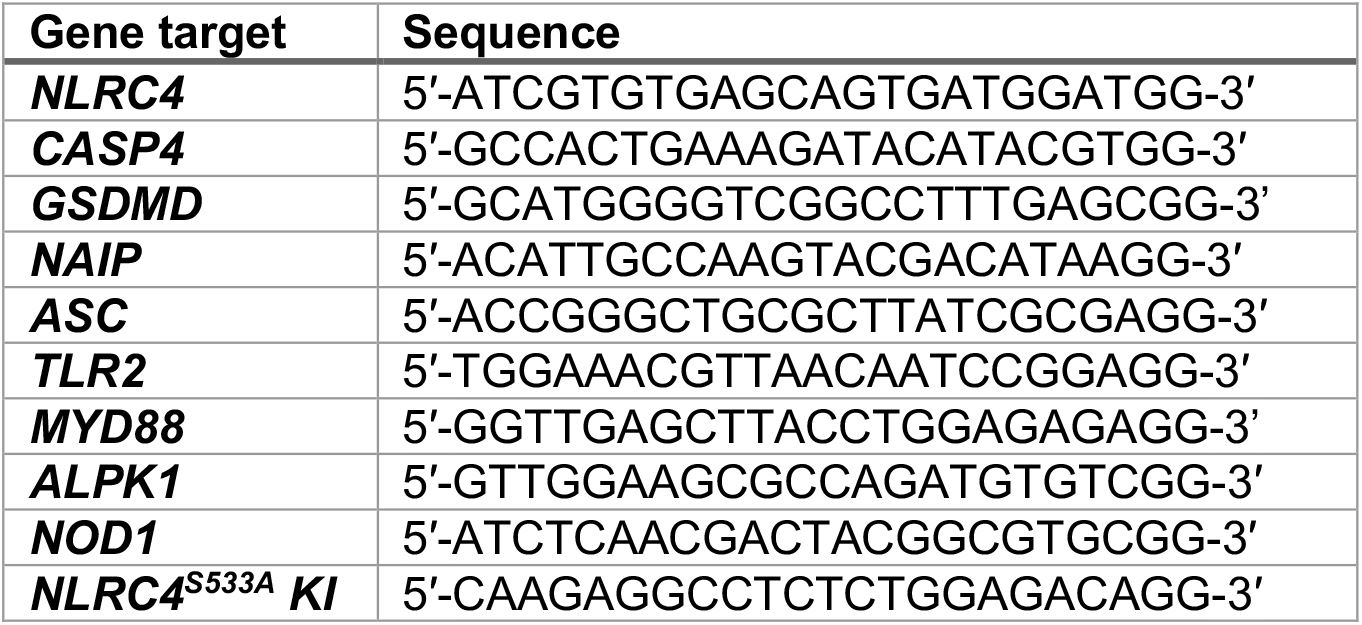
gRNA target sites used in this study.

**Table S3.** Primary antibodies used in this study.

| Target | Catalog | Dilution |
| --- | --- | --- |
| IL-1β | R+D, MAB201 | 1:1000 |
| vinculin | CST, #13901 | 1:1000 |
| phospho-p38 MAPK | CST, #9211 | 1:1000 |
| p38 MAPK | CST, #9212 | 1:1000 |
| phospho-ERK1/2 MAPK | CST, #4370 | 1:1000 |
| ERK1/2 MAPK | CST, #4695 | 1:1000 |
| HA | Roche, 11867423001 | 1:1000 |
| phospho-S533 (NLRC4) | ECM Biosciences, NP5411 | 1:1000 |
| β-actin | Santa Cruz Biotechnology, sc-47778 | 1:1000 |

## Supplemental Materials and Methods

### Immunoprecipitation

Treated cells expressing NLRC4-HA constructs were collected by cell lifter in cold PBS. Cells were lysed in 30 mM Tris-HCl, pH7.5, 150 mL NaCl with 0.5% Nonidet P40, with HALT protease and phosphatase inhibitors for 1 hr on ice. Lysates were centrifuged at max speed for 20 min. Supernatant was incubated with Pierce anti-HA magnetic beads for 3 hr at 4°C, rotating. After washing, samples were eluted in 1x Laemmli buffer at 70°C for 10 min. Samples were then used for Western Blot as described in methods.

### CRISPR-Cas9 Generation of THP-1 Knock-in Cell Line

For generation of knock-in NLRC4^S533A^, THP-1 cells were nucleofected with Alt-R™ S.p. Cas9 Nuclease V3 (IDT) complexed with sgRNA, and Alt-R™ Cas9 Electroporation Enhancer (IDT). Before combining with cells, IDT Alt-R™ HDR donor oligo was added to Cas9-sgRNA complexes. Cells were nucleofected in Lonza P3 buffer (Lonza, V4XP-3032) using the Lonza 4D Nucleofector Core Unit, program CM-137. Nucleofected cells were recovered in media containing 1 μM IDT Alt-R™ HDR Enhancer V2 (10007910) for 18 hr. Media was then changed to remove enhancer. Cells were plated for monoclones. Knock-in efficiency was sequenced and analyzed by ICE analysis (Edit Co). Guide and donor sequences were designed using Alt-R™ CRISPR HDR Design Tool (IDT). ssODN HDR oligo sequence: 5′-TCAGAATTTCTTGCTCAGTGGTGTTTTTCACACTTTGCAAGGCTTCCTGCCGCCAGAGAGGCCTCTTGGCGATGGAAAGTCCGAGAAGGCA-3′.

Guides used are listed in Table S2.

## Notes

### Competing Interest Statement

R.E.V. is on the scientific advisory boards of Tempest and X-biotix Therapeutics.

### Summary of Updates

Author affiliations updated; Figure 5 revised; updated and extended supplemental figures and tables.

